# Environment dependent benefits of inter-individual variation in honey bee recruitment

**DOI:** 10.1101/2021.08.18.456819

**Authors:** Supraja Rajagopal, Axel Brockmann, Ebi Antony George

## Abstract

Inter-individual differences in behaviour within the members of a social group can affect the group’s productivity. In eusocial insects, individual differences amongst workers in a colony play a central role division of labour and task allocation. Extensive empirical and theoretical work has highlighted variation in response thresholds as a proximate mechanism underlying individual behavioural differences and hence division of labour. However, other response parameters, like response probability and intensity, can affect these differences. In this study, we first extended a previously published agent-based model on honey bee foraging to understand the relative importance of response (dance) probability and response (dance) intensity in the task of recruitment. Comparing variation obtained from the simulations with previously published empirical data, we found that response intensity plays a more important role than probability in producing consistent inter-individual differences in recruitment behaviour. We then explored the benefits provided by this individual variation in recruitment behaviour to the colony’s collective foraging effort under different environmental conditions. Our results revealed that individual variation leads to a greater energetic yield per forager, but only when food is abundant. Our study highlights the need to consider all response parameters while studying division of labour and adds to the growing body of evidence linking individual variation in behavioural responses to the success of social groups.

## Introduction

Individuals from a population can vary in their behavioural decisions, even when responding to the same stimuli (Dall et al., 2012). Behavioural syndromes refer to the phenomenon in which individual variation in behavioural response is consistent over time and correlated across contexts (Sih et al., 2004). Consistent inter-individual differences in behaviour have been observed in a wide variety of taxa and diverse behavioural contexts, like aggression, courtship, and foraging behaviour (Bell et al., 2009). Differences in behavioural responses lead to behavioural specialisation amongst individuals and thereby influences how populations respond to perturbations (Wolf and Weissing, 2012). In eusocial insects, inter-individual differences in behaviour have been studied extensively in the context of behavioural specialisation associated with division of labour (Dall et al., 2012; Loftus et al., 2021).

Division of labour contributes to the phenomenal success of eusocial insect societies (Hölldobler and Wilson, 2009). Eusocial insect colonies consist of highly related workers who perform different tasks, like brood care and foraging, necessary for the survival and growth of the colony (Wilson and Hölldobler, 2005). Moreover, individuals performing the same task show differences in task performance, which can be consistent over time (Jeanson and Weidenmüller, 2013). Individual differences in task performance are linked to variation in the perception of the stimulus intensity as well as differences in the behavioural threshold of responding to the stimulus intensity (Beshers and Fewell, 2001). Theoretical studies on response thresholds suggest that variation in these thresholds is essential to maintain division of labour and ensure efficient utilisation of the workers in social insect colonies (Bonabeau et al., 1996; Johnson, 2010; Theraulaz et al., 1998). Empirical studies on response thresholds in a variety of tasks, like brood care and thermoregulation, have confirmed the link between thresholds and division of labour (Modlmeier et al., 2012; Weidenmüller, 2004).

Unfortunately, the focus on response thresholds has come at the cost of neglecting other response parameters that may be important for division of labour, like response probability and intensity (Cook et al., 2020; Jeanson and Weidenmüller, 2013; Ulrich et al., 2021).

An individual’s task performance for a given stimulus intensity is a combination of both its probability of responding to the stimulus and the intensity of its response, either in terms of the number or duration of response events (George and Brockmann, 2019; Weidenmüller, 2004). While thresholds determine only the minimum stimulus intensity at which individuals start responding, probability and intensity determine its response at a particular stimulus intensity. Individual variation in these response parameters can provide social insect colonies with additional flexibility in allocating their workforce and in responding to perturbations. Thus, a comprehensive understanding of the benefits of division of labour requires an exploration of these response parameters in addition to response thresholds. Despite their importance, empirical and theoretical studies on these response parameters in social insects are limited (Cook et al., 2020; Jeanson and Weidenmüller, 2013). An important requirement for such studies is a behavioural paradigm in which the responses of multiple individuals to the same stimulus can be quantified repeatedly.

The honey bee recruitment process consists of individual foragers, repeatedly visiting the same food source, using the highly stereotyped waggle dance behaviour to inform nestmates of the location and reward offered at the food source (von Frisch, 1967). An individual foragers motivation to recruit is dependent on both the reward offered by the food source as well as the colony’s nutritional state (De Marco and Farina, 2001; Nieh, 2010; Seeley, 1994; Thom, 2003). Foragers active at the same food source experience the same stimulus intensity, due to the same food reward and colony conditions, and hence the recruitment behaviour is an excellent paradigm to explore individual differences in behavioural responses. Recent work revealed that foragers show consistent inter-individual variation in their response probability (number of dances per trip) and intensity (average number of waggle phases per dance) of dance activity and, as a result, in their total recruitment activity for the same food source (George and Brockmann, 2019). While empirical studies have sought to understand how individual variation in recruitment behaviour benefits social insect colonies (Jones, 2004; Mattila and Seeley, 2010; Modlmeier et al., 2012), these are often difficult due to the need to control for multiple sources of individual variation (Jeanson and Weidenmüller, 2013; Ulrich et al., 2021). Simulations of the recruitment process can be a useful approach to understand how and under what conditions individual variation can potentially benefit social insect colonies, and thereby help design further behavioural experiments.

In this study, we explored how individual variation in recruitment activity affects the colony’s foraging behaviour by expanding upon a previously published agent-based model of the honey bee foraging process (Schürch and Grüter, 2014). We incorporated individual variation in response probability and intensity of the dance behaviour in the foraging process and perform two sets of experiments. In the first set of experiments, we estimated the relative importance of response probability and intensity in maintaining individual variation in the total recruitment activity of the agents. We did this by simulating a honey bee colony foraging under conditions mimicking experimental studies and validated the individual-specific parameters by comparing variation in the total recruitment activity in the simulations with empirical data (George and Brockmann, 2019). In the second set of experiments, we quantified the relative benefit of individual variation in recruitment activity under different environmental food availability conditions. By examining individual variation in recruitment behaviour from the perspective of both the cause of this variation in terms of response parameters and the consequences of this variation for the colony’s collective foraging activity, our study provides important insights into the role of inter-individual variation in social insect colonies.

## Methods

We simulated an agent-based model (ABM) of honey bee nectar foragers with inter-individual variation in the probability and intensity of their recruitment behaviour. Our model is built on top of an earlier ABM (Schürch and Grüter, 2014), which we modified to incorporate individual-level parameters. We used NETLOGO 5.3.1 to implement the model, and follow the “Overview, Design Concepts and Details” protocol to describe it (Grimm et al., 2006).

### Purpose

The overall purpose of the model was to explore the role of individual variation in the probability and intensity of recruitment behaviour amongst honey bee foragers in the colony’s foraging activity. We first quantified the effect of individual variation in probability and intensity in maintaining consistent inter-individual variation in dance activity for a particular food source. We calibrated the individual level parameter values by comparing variation in the total dance activity obtained from the simulations with empirical data (George and Brockmann, 2019). We then explored if individual variation in recruitment behaviour benefited the colony’s foraging efforts under different environmental conditions. Both probability and intensity (average number of waggle phases per dance) of dance activity is correlated with the forager’s perception of the food reward (Barron et al., 2007; George and Brockmann, 2019; Seeley, 1994, 1989). In our model, we implemented two individual state parameters: a probability modulator and an intensity modulator. The probability modulator regulates the probability that a honey bee performs a waggle dance on returning to the hive after foraging. The intensity modulator regulates the number of waggle phases that a bee makes in each waggle dance for a given nectar reward.

### State variables and scales

In all our model runs we simulated 300 foragers from one colony. The foragers were divided into two different types, scouts (n = 30) and recruits (n = 270), based on typical proportions of scouts and recruits in an *A. mellifera* colony (Seeley, 1995). We were interested in the consistency of recruitment activity to exploited food sources and hence focussed on the recruitment activity of the recruits. Scouts explore the environment for new food sources to exploit and usually abandon food sources even when they are still rewarding (Liang et al., 2012; Seeley, 1983). Recruits on the other hand get recruited to a food source (and can further recruit to the food source) and continue exploiting the food source till it becomes less rewarding (Biesmeijer and De Vries, 2001; Seeley, 1983). We were interested in individual variation in the recruitment activity of these recruits, as they bring in the bulk of the food necessary for the survival and growth of the colony. The states that individual agents could take in our model was the same as in the previously published model. Scouts could, at any time step, be in any of the 6 states (Fig. S1 *a*): idle in the colony, scouting for food sources, feeding at a food source, returning to the colony, recruiting of idle foragers to the newly discovered food source, returning to the nest without having discovered food. Recruits could be in any of the following 8 states (Fig. S1 *b*) at any time step: idle in the nest, waiting to be recruited, flying to a food source, feeding on a food source, returning to colony after feeding, recruiting new idle foragers, scouting for forage, returning to the colony without having discovered food. All agents were characterized by an identity number, a dance probability modulator, and a dance intensity modulator.

Our agents were located on a two-dimensional square grid with 201 × 201 patches. Following the previous model, the width of a patch corresponded to 100m in our model also. Thus, our agents could explore an area corresponding to a 20 km x 20 km square centred on the hive. Agents could occupy any position in this area and were not restricted to the edges of the patch. The agents’ hive was located at the centre patch and no food sources were present in this patch throughout the simulations.

Two different sets of experiments were performed which differed in the number and persistence of the food availability in the environment. In experiment 1, we simulated a foraging environment for our agents with no temporal change in the distribution of nectar sources for a short duration of 5 days. This was done to ensure that we could easily compare our simulation results with previous experimental data (George and Brockmann, 2019). In experiment 2, we simulated a more natural foraging environment with multiple transient food sources for a longer duration of 51 days. Similar to the earlier published model (Schürch and Grüter, 2014), we tested three different environmental conditions in this experiment: low food density, medium food density and high food density.

Simulations were run in discrete time-steps. Each time step corresponded to 10 seconds in real-time and each day consisted of 5000 steps, with agents active up to the 4000^th^ time step on each day (corresponding to 11.11 hours of real time). The weather was not simulated, and so the general foraging activity of the colony remained similar throughout the day.

### Process overview and scheduling

Each simulation was run for 5 days (experiment 1) or 51 days (experiment 2), consisting of a short non-experimental phase at the beginning (2 days for experiment 1 and 3 days for experiment 2). The non-experimental phase allowed foragers to learn about the location of food sources around their colony. Scouts and recruits followed similar processes while foraging (Fig. S1 a and b). The recruitment process in our model was based on the earlier model. After unloading the nectar, individual agents could dance to recruit one other agent to the food source, with the probability of recruitment dependent on the duration of the dance.

The probability that a bee will dance on returning to the hive was determined by her probability modulator, the nectar quality in terms of sugar concentration (Tautz and Sandeman, 2003; Waddington, 1982), the distance of the nectar source from the hive (Seeley, 1994) and the current influx of returning foragers (Seeley, 1989). The quality of the food sources and their distances from the hive was kept the same throughout, and the current influx of returning foragers was the same for bees dancing at the same time step. Thus, in our model, individual differences in dance probability for foragers visiting the same food source were a direct result of individual differences in the probability modulator. If a bee decides to dance, the intensity of her dance is determined by nectar source distance, nectar quality (Seeley, 1994) and her dance modulator. Individual differences in dance intensity for foragers visiting the same food source were a direct result of individual differences in the intensity modulator.

Scouts and recruits became idle after dancing. Agents in an idle state could leave the nest with probability *p*_*exit*_. Scouts constantly searched for new food sources, while recruits showed high patch fidelity (Al Toufailia et al., 2013; Moore and Doherty, 2009). Each day recruits, that had previously foraged, decided to either continue foraging or abandon the patch. The probability to abandon the patch depended on the individual probability modulator, nectar quality and nectar source distance.

#### Experiment 1: Individual variation in probability and intensity and consistent differences in recruitment behaviour

We quantified how much individual variation in probability and intensity of dancing contributes to consistent inter-individual differences in the total dance activity of individual foragers. To measure the consistency of dance activity, we ensured that foragers had access to four food sources of the same quality (same sucrose concentration at the same distance from the hive in the four cardinal directions). The food source was available throughout the day, and we quantified the dance activity of foragers at this food source for 5 days (2 days to acclimatize followed by a 3-day experimental phase). We quantified individual differences in the dance activity in the experimental phase and obtained the repeatability estimates of the total dance activity. This was then compared to empirical repeatability estimates obtained from observations of groups of foragers trained to an ad-libitum food source over 3 days (George and Brockmann, 2019). The comparison allowed us to calibrate the variation in probability and intensity modulators to obtain empirical levels of variation in recruitment activity and further quantify the relative importance of probability and intensity in maintaining consistent differences in recruitment activity (through a sensitivity analysis).

#### Experiment 2: Individual variation in recruitment behaviour and colony foraging activity

Environmental conditions can affect the benefits of information transfer through the recruitment process in social insects, with the greatest benefit often seen in low food density conditions (Dornhaus et al., 2006; Goy et al., 2021; Schürch and Grüter, 2014). In this experiment, we explored how inter-individual variation in the recruitment behaviour further benefits the colony’s foraging activity. We hypothesise that the benefits of inter-individual variation in recruitment behaviour will be dependent on the environmental conditions. Inter-individual variation in the recruitment behaviour was implemented as a function of individual variation in both response probability and intensity, which we obtained from the previous experiment. We tested the colony’s foraging activity under three different environmental conditions: low, medium, and high food density (*d*_*patch*_ corresponding to 0.01, 0.05 and 0.1 respectively).

In both experiments 1 and 2, we tested four different variations of our model:

Model 1: Honey bee forager agents were all similar with no variation in individual state variables of probability modulator (*I*_*p*_) and intensity modulator (*I*_*i*_).

Model 2: Every agent was characterised by a different value of *I*_*p*_ and the same value of *I*_*i*_. Honey bee foragers showed inter-individual differences in their probability of dancing for any given food reward. The probability modulator for each agent was obtained from a normal distribution [Fig. S2 a; mean ± standard deviation (sd) of *I*_*p*_ = 0.255 ± 0.1].

Model 3: Every agent was characterised by a different value of *I*_*i*_ and the same value of *I*_*p*_. Honey bee foragers showed inter-individual differences in their intensity of dances for any given food reward. The intensity modulator for each agent was obtained from a normal distribution (Fig. S3 a; mean ± sd of *I*_*i*_ = 0.255 ± 0.1).

Model 4: Each agent was characterized by a different value of *I*_*p*_ and *I*_*i*_, both of which were obtained from normal distributions (mean ± sd = 0.255 ± 0.1). Honey bee foragers showed inter-individual differences in both the probability and intensity of dancing for any given food reward.

An agent with the mean values of *I*_*p*_ and *I*_*i*_ in models 2, 3 and 4 was similar to all agents in model 1.

### Design Concepts

In our model, individual honey bee foragers collected food from various food sources offering different sucrose concentrations present throughout the environment around the hive which was centrally located. The amount of food brought back is equivalent to the energy procured by the colony and thus its fitness.

#### Environmental Stimuli

The honey bee foragers could evaluate whether the time of day was suitable for foraging and the quality of the food sources. The probability and intensity of dancing depended on this quality. Additionally, the probability of recruits to persist at the food source also depended on this quality.

#### Biotic stimuli

Scouts and recruits could pass on the information about nectar source location to other recruits. Recruits mostly flew out only after receiving this information.

#### Stochasticity

The probability and intensity modulators were assigned stochastically. They were picked randomly from a normal distribution and assigned to the agents. Agent decisions regarding leaving the hive, dancing for a food source, learning the location of a nectar source and abandoning a nectar source were all probabilistically determined.

#### Observation

Different emergent properties of the model were observed in experiments 1 and 2. In experiment 1, the food patch visited and the number of dance circuits in each dance was recorded and used to obtain the total number of dance circuits made by each agent for each day. In experiment 2, the mean energy yield per forager, the median foraging distance, and the mean search time in addition to the mean recruitment activity for each day were quantified. The former three properties represent mutually non-exclusive mechanisms by which inter-individual variation in recruitment can benefit the colony’s foraging effort: through an increase in the energy brought in by foragers, a reduction in the distance flown by foragers to find a rewarding food source and a reduction in the time spent by foragers searching for food sources independently (because of an increased probability of recruitment to a rewarding food source).

### Initialisation

All agents were idle at the start of the simulation. Environmental food availability at the start of the model depended on the experiment. In experiment 1, four equally rewarding (0.8 M sucrose concentration) nectar sources were available in the environment at a distance of 80 patches from the hive. The four patches were distributed around the nest with a 90° angle between consecutive patches. These patches never got exhausted for the duration of the simulations (5 days), similar to artificial feeders used in behavioural experiments with honey bees (Seeley, 1995). In experiment 2, multiple food sources of varying quality were available in the environment around the hive at the beginning of each simulation as initialised in the previously published model (Schürch and Grüter, 2014). The number of food sources depended on the environmental food density (*d*_*patch*_). Each patch in the simulated world was randomly allocated as a foraging patch or not with a probability equal to *d*_*patch*_. The quality of the food source was a random value from a normal distribution with a mean of *q*_*patch*_ and a standard deviation of *σ*_*patch*_. All food sources were finite with a maximum age (*a*_*max*_) of 14 days. The current age of the patch at the beginning of the simulation was a random value between 1 and 14 with equal probability.

### Sub models

#### The Lévy flight

The movement of agents during a search followed an optimal Lévy flight pattern (Reynolds et al., 2009). This sub model was obtained from Schürch and Grüter, 2014. Agents moved in bouts after initially choosing a random direction. Before the start of the bout, a total travel distance for the bout was calculated using the formula:

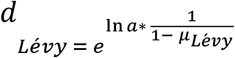

Here, *a* was obtained randomly from a uniform distribution bound by [0,1). The value for *µ*_*Lévy*_ was kept at 2.4 by default (Reynolds et al., 2009). Once the distance of the bout was calculated, agents would then move in the chosen direction in subsequent time steps, till they had reached the end of the distance. Then, the whole process was repeated till the agent found a food source.

#### Individual decision to dance for food sources

On returning to the hive, successful scouts and recruits would first unload the nectar that they had brought into the hive. They could then communicate information about the location of the food source to their nest-mates by performing a waggle dance. The forager would decide to dance only after the agent has finished unloading the nectar brought back from foraging. The equation that linked the probability of dancing to the food reward was

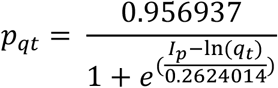

Here *I*_*p*_ was the individual probability modulator and *q*_*t*_ was the energetic value of a food source, given by

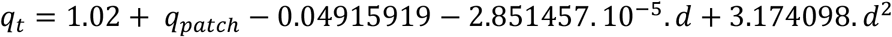

where *q*_*patch*_ was the sucrose concentration of the food source and *d* was the distance of the food source from the hive.

Our equation was modified from the earlier model, which in turn used data from previous empirical studies (Boch, 1956; Seeley et al., 1991). We incorporated individual probability modulators, thereby giving each individual agent a different quantitative relationship between food quality and dance probability. The mean value for the probability modulator was obtained based on two assumptions: 1) all individual agents had the same qualitative relationship between patch quality and dance probability as described in Schürch and Grüter, 2014 and 2) the probability modulators of the population of agents were normally distributed. We chose the mean and standard deviation of the probability modulator based on the average value of the curves fit for each agent, such that our two assumptions were met.

The dance probability also depended on the influx of other foragers,

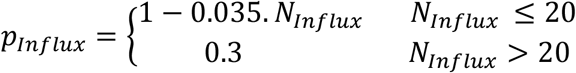

Thus, the final dance probability was given by

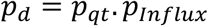

#### Individual decision about the intensity of dances

Dance intensity is the number of waggle phases that a bee makes per dance. The relationship between food quality and the number of waggle phases is linear (Seeley, 1994). In our model, each bee was assigned a quantitatively different slope that predicted how much they would dance after returning to the hive.

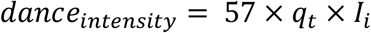

Here *I*_*i*_ referred to the individual intensity modulator and *q*_*t*_ was the energetic value of a food source.

To obtain the value of the intensity modulator, we made 4 assumptions: 1) all individual agents had the same qualitative linear relationship between patch quality and dance intensity as observed in Seeley, 1994, 2) the intensity modulators of the population of agents was normally distributed, 3) all agents had the same intercept of zero for the linear link between patch quality and dance intensity, i.e., there was no dance activity for a patch of zero quality and 4) there is an upper limit to the number of waggle phases for the most rewarding food source due to energetic constraints (Seeley, 1995). Based on these assumptions, we fit individual curves for each agent and chose the mean and standard deviation of the intensity modulator.

The intensity of the dance determines the duration of the dance. In the previously published model, all dances had the same intensity and lasted for 10 seconds (equal to the minimum duration for an agent to change its state). In our model, the more intense the dance, the longer it lasted. Each dance circuit lasted for around 3.33 seconds such that within each discrete time-step in the model, an agent could complete 3 dance circuits. As a result, agents with steeper intensity curves (higher *I*_*i*_ values) spent more time performing the recruitment behaviour. This had a corresponding effect on the agents recruited through the waggle dance. In the previous model (Schürch and Grüter, 2014), each dance had a 25% chance to recruit a forager, in accordance with empirical data (Dornhaus et al., 2006; Gould et al., 1970; Seeley and Towne, 1992; Seeley and Visscher, 1988; Tautz, 1996). In our model, we adjust the recruitment process so that every 10 dance circuits had a 25% chance to recruit a forager. Thus, longer dances probabilistically recruited more agents.

#### Individual decision to abandon food sources

The probability of abandoning the nectar source was modelled analogously to the probability of dancing (Schürch and Grüter, 2014; Seeley, 1989) with the probability modulator playing a role in the decision to abandon a food source:

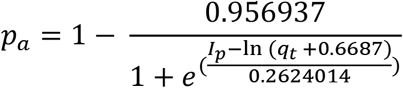

Thus, there was a correlation between an individual’s decision to dance for a food source and the decision to abandon the food source, tied to the individual’s perception of the food reward (George and Brockmann, 2019; Seeley, 1989).

### Sensitivity Analysis

We ran each experimental simulation multiple times to account for the stochasticity in the models. In experiment 1, each of the 4 variations of the model was run 100 times. In experiment 2, each of the 12 combinations of model and food density variation (4 variations of the model and 3 different food densities) were run 40 times.

Further, to test how sensitive our model was to changes in our two individual-level response parameters, we ran a sensitivity analysis based on model 4 in experiment 1. We chose this experiment as the controlled food availability allowed us to check the relative effect of probability and intensity on consistent inter-individual variation in recruitment activity. We chose this model as it incorporates individual variation in probability and intensity of recruitment, unlike the other 3 models. We varied the distribution of the probability (Fig. S2 b) and intensity modulators (Fig. S3 b), to change the extent of inter-individual differences in probability and intensity. The probability equation was modified as shown below to vary the spread of the individual modulators:

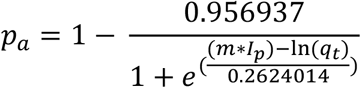

where m took a value in between 0.25 and 2 in steps of 0.25. To change the spread of the individual variation in intensity, we changed the standard deviation of the normal distribution of *I*_*i*_, while keeping the mean constant. The standard deviation was changed from 0.01 to 0.1 in steps of 0.01. We then quantified the consistency in inter-individual variation in the total recruitment activity for each of the 80 combinations (8 levels of variation in *I*_*p*_ and 10 levels of variation in *I*_*i*_) and ran 10 simulations of each version for a total of 800 simulations. We varied the spread of the two modulators in this way to keep the coefficient of variation of both modulators similar. This allowed us to directly compare the effect of variation in probability and intensity in maintaining consistent inter-individual differences in recruitment behaviour.

Sensitivity analyses regarding other parameters in the model, like the colony size, probability of scouts to exit the nest *p*_*exit*_, Levi-flight parameter *µ*_*Lévy*_ and energy costs of recruiting were explored in the previous study (Schürch and Grüter, 2014). The model used in their analysis was analogous to model 1 from experiment 2 in this study with medium food density.

### Statistical Analysis

#### Experiment 1

In this experiment, we were interested in quantifying the effect of individual variation in probability and intensity of dance activity on maintaining consistent inter-individual differences in recruitment behaviour. To do this, we first shortlisted foragers who consistently visited the same patch over the three days to minimise any variation in dance activity due to a forager switching states. As scouts abandoned food sources readily, they were automatically removed in our analysis. We then used generalized linear mixed-effects models (GLMMs) with the total dance activity in a day as the response variable and individual agent identity as the random effect and a Poisson error distribution to quantify the repeatability of recruitment behaviour. Repeatability estimates provide a quantification of the consistency in inter-individual differences for a behaviour and are based on a comparison of intra-individual variation to inter-individual variation (Nakagawa and Schielzeth, 2010). We obtained a single repeatability estimate for each of the 100 simulations runs of each experiment. The repeatability estimates were then compared using a beta regression model with the estimate as the response variable and the model as the predictor (categorical variable of 4 levels, corresponding to models 1, 2, 3 and 4). A beta regression model was used as the values of repeatability estimates was a proportion between 0 and 1. We compared the 4 different models against each other and corrected for multiple comparisons using Tukey’s HSD.

#### Experiment 2

In this experiment, we were interested in quantifying the effect of individual variation in recruitment behaviour on the colony’s collective foraging activity. We observed 4 different output parameters on each day for all the simulation runs: mean dance time per forager, mean yield per forager, median foraging distance and mean search time per forager. We then compared these parameters across the 4 different models and the 3 different environmental food density conditions using linear mixed-effects models (LMMs). We built separate LMMs for each parameter, with the parameter of interest as the response variable, an interaction between the food density (categorical variable of 3 levels) and the model (categorical variable of 4 levels) as the predictor and day nested within simulation number as the random effect. We compared the effect of the different models against each other within each level of food density and corrected for multiple comparisons using Tukey’s HSD.

#### Sensitivity Analysis

For the sensitivity analysis, we first obtained the repeatability estimates for all 800 simulation runs as in experiment 1. Then, a LOESS model was fit to this data with the mean repeatability estimate for each of the 80 combinations of probability and intensity modulator as the response variable and an interaction between the coefficient of variation in the probability modulator and the coefficient of variation in the intensity modulator as the predictor variable. We then predicted the repeatability estimates for other combinations of variation in the probability and intensity modulator from this fitted model. Finally, contour plots were used to visually examine the effect of the variation in probability and intensity modulators on the repeatability of dance activity.

We built and ran the models in NETLOGO 5.3.1 (Wilensky, 1999). The model outputs were cleaned up using the pandas package (McKinney and others, 2010) in Python (Van Rossum and Drake, 2009). We used R software (R Core Team, 2018) along with the RStudio IDE (RStudio Team, 2016) to perform all the statistical analyses. We built the GLMMs using the rptR package in R (Stoffel et al., 2017). For the beta regression, we used the betareg package (Cribari-Neto and Zeileis, 2010), for LMMs we used the glmmTMB package (Brooks et al., 2017) and for multiple comparisons, we used the emmeans package (Lenth, 2019). We fit the LOESS model using the base package in R. We validated the model assumptions for each of the mixed-effects models using visual inspections of residual and normality plots through the DHARMa package (Hartig, 2018). For data visualisation, we used the ggplot2 (Wickham, 2016) and cowplot (Claus O. Wilke, 2018) packages in R. All the code produced as part of this manuscript can be found at the Zenodo repository (George et al., 2021).

## Results

### Experiment 1: Individual variation in probability and intensity and consistent differences in recruitment behaviour

Incorporating individual variation in probability and intensity increased the consistency of inter-individual differences in recruitment activity amongst the agents in the simulations (Table S2 and Fig. 1a, models 2, 3 and 4 all had significantly higher repeatability estimates compared to model 1 at the p < 0.001 level). However, repeatability estimates of the total recruitment activity were much higher in model 3 which only incorporated individual differences in intensity, as compared to model 2 which only incorporated individual differences in probability (Table S2 and Fig. 1a, model 3 - model 2: repeatability estimate difference = 0.524, z-ratio = 82.911, p < 0.001). Further model 4, which incorporated both individual variation in probability and intensity, had lower repeatability estimates compared to model 3, which had only individual differences in intensity (Table S2 and Fig. 1a, model 3 - model 4: repeatability estimate difference = 0.139, z-ratio = 18.152, p < 0.001). Both model 3 and model 4 had mean repeatability estimates comparable to empirical observations (Fig. 1b). However, estimates obtained from model 4 were the closest to empirically obtained repeatability estimates.

**Figure 1:**
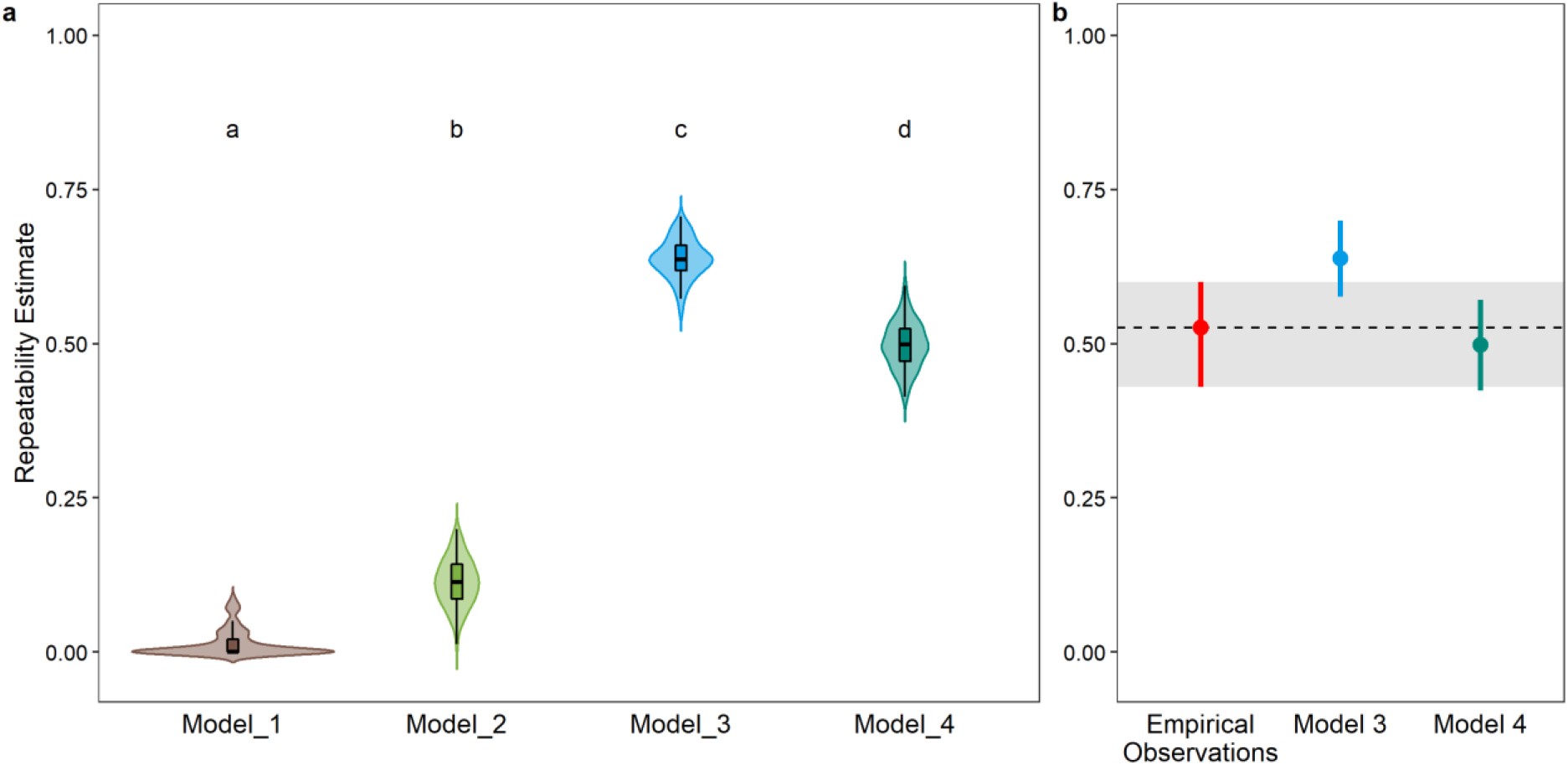
Repeatability estimates of recruitment behaviour (i.e., total dance activity) of agents in the model. (a) Violin plots of the distribution of repeatability estimates from 1200 simulation runs of the four different models. Inset boxplots show the median (central black line), the first and third quartiles (lower and upper edges of the box) and 1.5 times the interquartile range between the first and third quartile (whiskers). Alphabets above each violin plot represent the results of the post-hoc pair-wise comparison of repeatability estimates between the different models, with different alphabets representing significant differences at the p < 0.05 level. (b) Comparison of the repeatability estimates obtained from models 3 and 4 with estimates obtained from empirical observations (data in George and Brockmann, 2019). The circles represent the repeatability estimate (mean of the estimates in the case of the models) and the error bars represent the 95% confidence interval. The grey shaded region represents the 95% confidence interval of the repeatability estimate from empirical observations.

Our sensitivity analysis corroborated results from experiment 1. Individual variation in intensity had a stronger effect on maintaining consistent inter-individual differences in dance activity at the colony level as compared to a similar level of individual variation in probability (Fig. 2).

**Figure 2:**
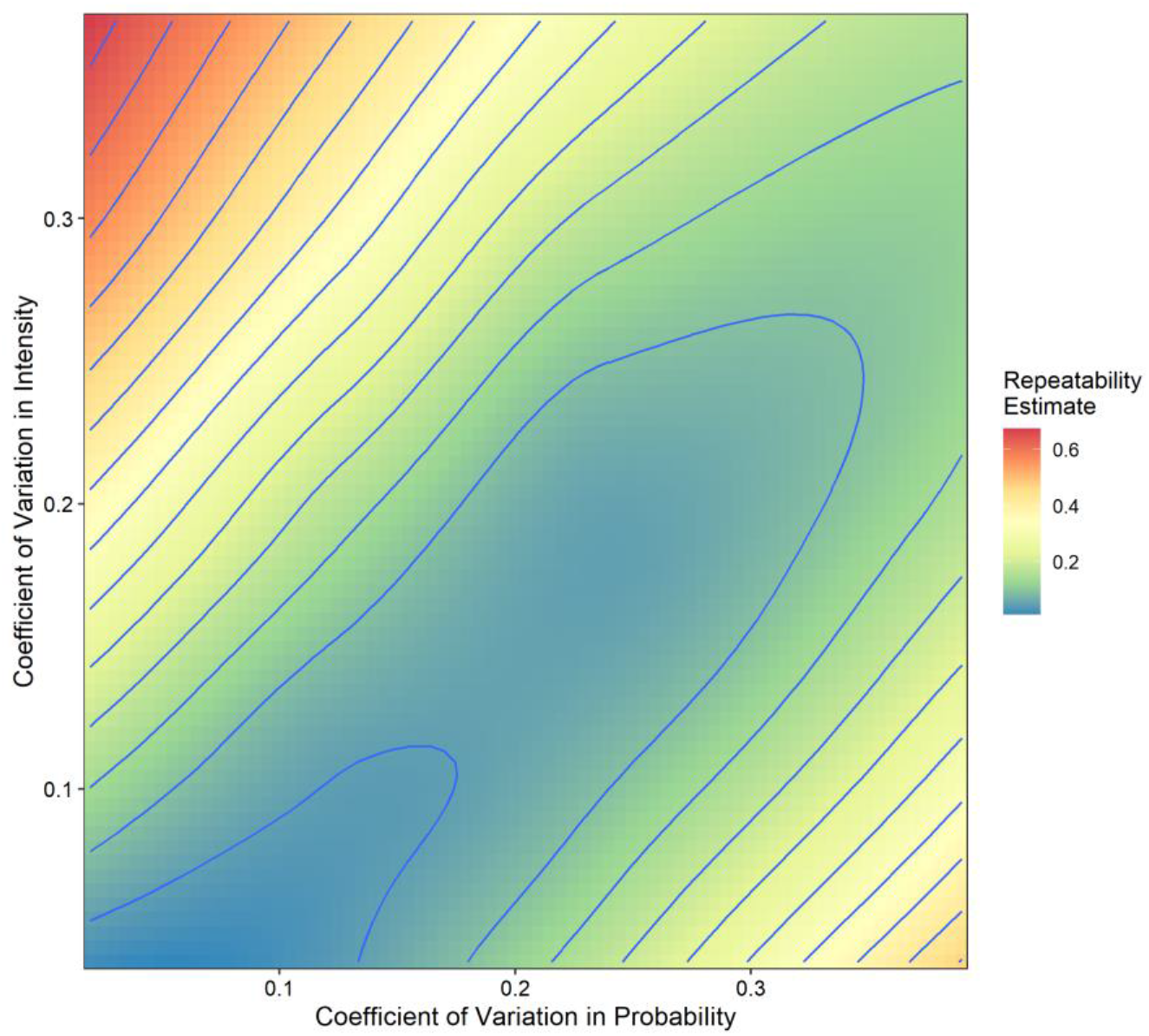
Contour plot of the sensitivity analysis on the effect of individual variation in probability and intensity of dancing on the repeatability estimates of recruitment behaviour. The contour plot is coloured based on the value of the repeatability estimate predicted for each combination of coefficient of variation in probability and intensity.

### Experiment 2: Individual variation in recruitment behaviour and colony foraging activity

Incorporating inter-individual variation in recruitment behaviour affected the colony’s foraging dynamics in an environmental dependent manner, with the strongest effects at medium and high food densities (Table S3 and Fig. 3). Inter-individual variation in recruitment behaviour led to significantly shorter recruitment time at the group level under all food density conditions (Fig. 3c, model 1 – model 4: low food density – difference estimate = 3, t-ratio = 10.08, p < 0.001; medium food density – difference estimate = 2.32, t-ratio = 7.77, p < 0.001; high food density – difference estimate = 2.27, t-ratio = 7.63, p = < 0.001). There was a similar trend in the opposite direction in the mean yield with individual variation in recruitment behaviour leading to higher yield at medium and high food densities (Fig. 3d, model 1 – model 4: low food density – difference estimate = 0.64, t-ratio = 0.74, p = 0.881; medium food density – difference estimate = -3.08, t-ratio = -3.55, p = 0.002; high food density – difference estimate = -3.44, t-ratio = -3.95, p < 0.001).

**Figure 3:**
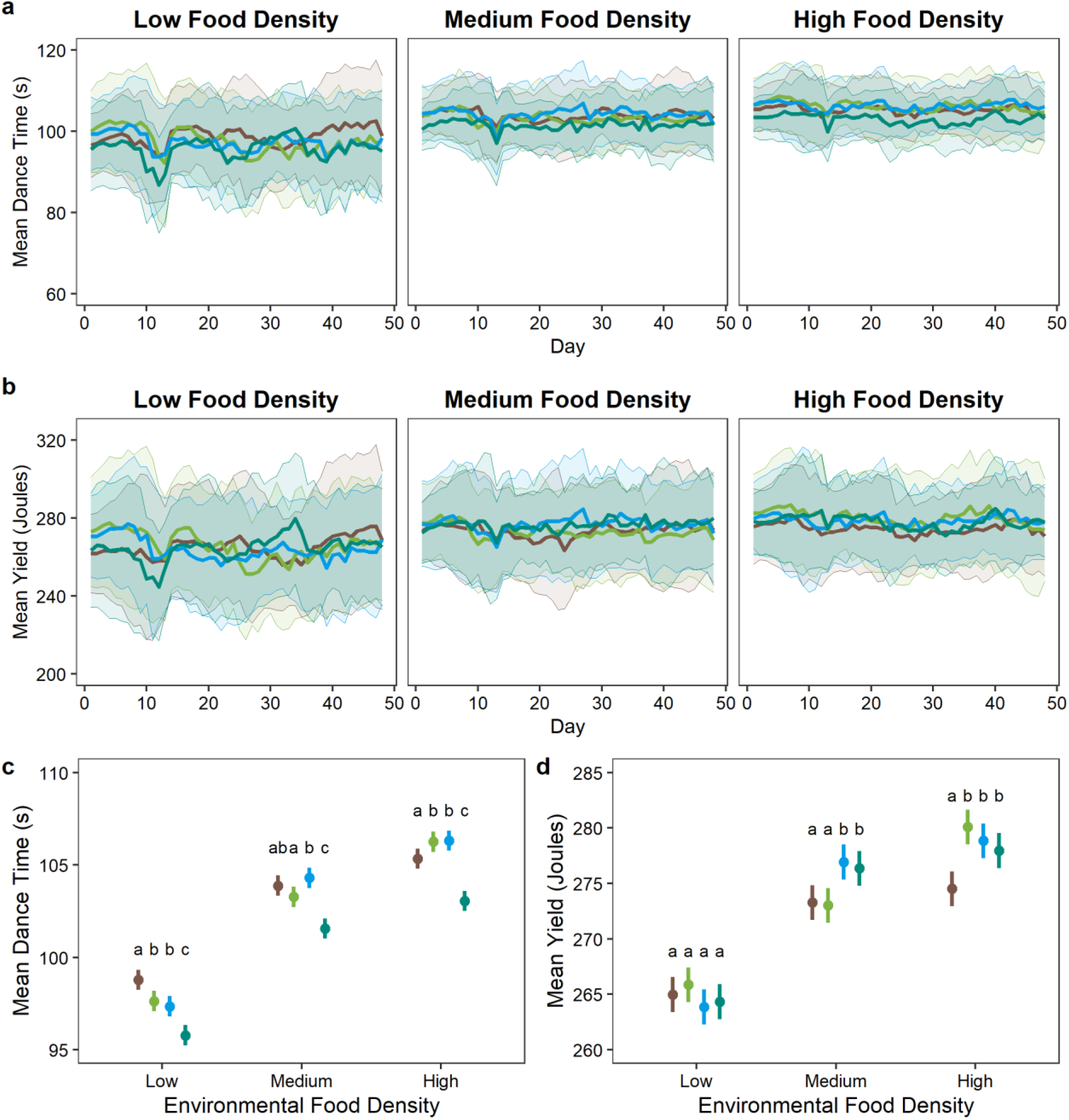
Effect of individual variation in recruitment behaviour on the colony’s foraging activity under different environmental conditions. (a and b) Line plots showing the change in (a) mean dance time per forager in seconds and (b) mean yield per forager in joules throughout the simulations. The central line represents the mean value of all agents on each day from 40 simulations while the shaded region represents the standard deviation around this mean. (c and d) Comparison of the (c) mean dance time and (d) mean yield per forager, averaged over the whole simulation duration for all 40 simulations, across models and food density conditions. Circles represent the mean and error bars represent the standard deviation. The lines and shaded region in (a) and (b) and the circles and error bars in (c) and (d) are coloured according to the model: model 1 – brown, model 2 – green, model 3 – blue, model 4 – teal.

There was no significant difference in the median foraging distance between the model with and without individual variation in probability and intensity in any of the food density conditions (Fig. S4, model 1 – model 4: low food density – difference estimate = 0.001, t-ratio = 0.08, p > 0.999; medium food density – difference estimate = -0.003, t-ratio = -0.2, p = 0.997; high food density – difference estimate = -0.009, t-ratio = -0.66, p = 0.912). The mean search time per forager in the model with and without individual variation in recruitment activity followed the same pattern as the median foraging distance, with no difference in any of the food density conditions (Fig. S4, model 1 – model 4: low food density – difference estimate = -0.54, t-ratio = -0.22, p = 0.996; medium food density – difference estimate = -0.001, t-ratio = -0.0005, p > 0.999; high food density – difference estimate = 1.16, t-ratio = 0.48, p = 0.964).

Incorporating individual variation in only one of the two response parameters did not produce the same decrease in the mean time spent dancing per forager as incorporating individual variation in both parameters did (Table S2 and Fig. 3, comparison of model 4 versus models 2 and 3). On the other hand, it produced similar levels of mean yield, especially in the high food density conditions. Further, the median foraging distance and mean search time was comparable across all models in the medium and high food density conditions, with a slight increase in the median foraging distance and mean search time in the low food density condition (Table S2 and Fig. S4).

## Discussion

Our simulations revealed two important aspects of individual variation in recruitment activity amongst honey bees. First, variation in response intensity contributed more to producing consistent variation in recruitment activity as compared to variation in response probability. At the same time, incorporating individual variation in both response parameters was necessary to recapture empirically observed levels of consistent inter-individual variation in recruitment activity. Second, individual variation in recruitment activity led to a decrease in the average recruitment activity of the colony and benefitted the colony’s foraging activity by increasing the mean yield per forager. However, this benefit of individual variation in recruitment activity was only apparent at medium and high food density conditions, and not at low food density conditions. By focussing on the response parameters responsible for individual variation in task performance, our study adds to the growing body of literature highlighting the importance of response probability and intensity in addition to response thresholds in social insect division of labour.

The three response parameters may play different roles in modulating individual differences in task performance. Empirical work on individual variation in honey bee recruitment behaviour showed that response intensity is weakly correlated with gustatory response thresholds whereas response probability is not (George et al., 2019). Similarly, a study on variation in recruitment behaviour at the level of patrilines revealed no correlation between thresholds and probability (Mattila and Seeley, 2010). Our simulation results now reveal that although individual variation in both response probability and intensity is necessary to obtain empirical levels of inter-individual differences in recruitment activity, the latter is more important. At the same time, there is evidence that response probability may play a vital role in the social modulation of recruitment. Social interactions between foragers and receivers who unload the nectar inform foragers of the nutritional state of the colony and can change their recruitment behaviour (Cook et al., 2020; De Marco and Farina, 2001; Seeley, 1989). We could not simulate these interactions in greater detail due to the different spatial scales of these interactions (within the hive, over centimetres) and the foraging behaviour (outside the hive, over kilometres). However, in a previous experiment, manipulating the nectar influx, and thereby changing these social interactions of foragers (Seeley, 1989), affected the probability but not the intensity of the recruitment behaviour (George and Brockmann, 2019). Taken together, these results imply that response intensity and threshold is likely linked to an internal behavioural state responsible for innate variation in the recruitment activity of individual foragers. Modulation of response probability via social interactions allows the colony further flexibility in utilising this basal level of inter-individual variation.

Variation in response thresholds is generally considered to be the major physiological basis underlying task specialisation and division of labour in social insects (Beshers and Fewell, 2001; Page and Mitchell, 1998). The simplest models on division of labour assume that individuals have fixed response thresholds, whereas more complex models involve flexible individual thresholds which are modulated by factors like task performance and social interactions (Theraulaz et al., 1998). In contrast, a few empirical studies have indicated that individual differences in task performance within social insect colonies are often based on an intricate relationship between thresholds and two other response parameters: response probability and response intensity (Cook et al., 2020; Jeanson and Weidenmüller, 2013; Ulrich et al., 2021; Weidenmüller, 2004). The relative importance of the response parameters in task performance may vary depending on the task and the social context. In bumblebees, for example, a combination of response probability and intensity was most predictive of individual variation in task performance during thermoregulation (Garrison et al., 2018). Response thresholds did not predict individual task performance and differed in the same individuals when they were observed singly and in social groups. In honey bee recruitment, response probability is sensitive to social signals, whereas response intensity remains unaffected (George and Brockmann, 2019). Differences in the role of probability and intensity were also apparent in our simulations, and a useful direction for future simulations would be to incorporate individual variation in thresholds into our model.

Individual variation in recruitment behaviour can provide additional benefits to a social insect colony on top of those provided by recruitment itself. Our results reveal potential trade-offs during the collective foraging process in honey bees, specifically between recruitment to and exploitation of a food source. Inter-individual differences in the waggle dance would allow some foragers to recruit more foragers to a newly discovered food source while others can continue exploiting it. This division of labour within groups of foragers active at the same food source leads to an overall decrease in the average recruitment activity in our simulations. Individual differences in recruitment should also ensure that recruitment is further biased towards highly profitable food sources (Seeley, 1994, 1989, 1986). The high reward offered by these food sources should also lead to continuous exploitation and lowered abandonment by foragers. In line with this, there was an increase in the colony’s productivity in models incorporating individual variation in recruitment. At the same time, the benefits of individual variation in recruitment are likely to be seen after the discovery of food sources and not before. Consistent with this expectation, individual variation in recruitment did not affect the median foraging distance of the foragers or the average time spent searching for food sources by the foragers in our simulations.

The benefits of spatially directed recruitment, e.g., through the waggle dance, is more apparent under heterogeneous environmental conditions with low food densities (Bailis et al., 2010; Beekman and Lew, 2007; Dornhaus et al., 2006; Goy et al., 2021; Schürch and Grüter, 2014, but see I’Anson Price et al., 2019). When food is scarce, a decrease in recruitment activity should lead to a decrease in the colony’s productivity due to lower number of foragers being directed to the most profitable food sources. On the other hand, when food sources are plentiful, the energy and time costs of recruitment would likely outweigh the benefits provided by this behaviour (Dornhaus et al., 2006; I’Anson Price and Grüter, 2015; Schürch and Grüter, 2014). Our simulations reveal an under-appreciated role of inter-individual differences in spatially directed recruitment at high food density conditions. When food is plentiful, individual differences lead to a lowered average recruitment activity as well as a corresponding increase in the energy gain for the colony. Thus, these differences can offset the energy and time costs of recruitment and ensure that recruitment is still beneficial to the colony under such environmental conditions. One important aspect of most studies on the benefits of the waggle dance, including the current study, is that they are limited to *A. mellifera*. The other extant honey bee species show marked differences in their behaviour and ecology in comparison with *A. mellifera* (I’Anson Price and Grüter, 2015), particularly with respect to the waggle dance (Beekman et al., 2015; E. A. George et al., 2021; Kohl et al., 2020). Species in temperate regions (e.g., *A. mellifera* and some populations of *A. cerana*) have a shorter time window for foraging throughout the year compared to species in tropical regions (e.g., *A. florea, A. dorsata* and *A. cerana*) but must establish large winter food stores (Ji et al., 2020). The role of individual variation under these different environmental conditions is unknown and comparative studies on individual variation within the genus *Apis* would be an important avenue for future research.

Individual variation in social groups is used as a bet-hedging strategy to buffer against environmental variation (Honegger and de Bivort, 2018; Hopper, 1999). However, the perspectives behind the studies on this variation have differed between eusocial groups and other social groups (Dall et al., 2012). In eusocial insects, theoretical and empirical studies across species (honey bees, bumblebees, and ants) and tasks (brood care, thermoregulation, foraging and recruitment) have focussed on the role played by individual variation in task allocation (Beshers and Fewell, 2001; Jeanson and Weidenmüller, 2013; Mattila and Seeley, 2010; Modlmeier et al., 2012; Weidenmüller et al., 2019; Westhus et al., 2013; Wiernasz et al., 2008). On the other hand, in social groups, individual variation has been studied in terms of the range of behavioural responses a group has access to, which can, in turn, affect its productivity, e.g., in the case of foraging in birds and fishes (Aplin et al., 2014; Dyer et al., 2009; Wolf and Weissing, 2012). Irrespective of these different perspectives, individual variation in behavioural responses ultimately affects group success across social groups. Intriguingly, a new theoretical framework proposes to integrate concepts from task allocation in eusocial insects and animal personality studies in social groups (Loftus et al., 2021). A careful exploration of such a framework (Pinter-Wollman, 2021) would enable a better understanding of the link between consistent inter-individual differences in behavioural responses and division of labour in social insects.

In conclusion, our study reveals both the relative importance of two response parameters in producing consistent inter-individual differences in task performance during recruitment and the effect of these differences on a honey bee colony’s foraging activity. These results corroborate results from previous experimental and theoretical studies on the benefits of inter-individual variation for collective task performance in animal groups including social insects (Aplin et al., 2014; Michelena et al., 2010; Seeley, 1994). They also highlight the role played by individual differences in offsetting the costs of recruitment at the group level under different environmental conditions (Beekman and Lew, 2007; Dornhaus et al., 2006; Schürch and Grüter, 2014). Importantly, our study adds to the accumulating evidence on the importance of multiple response parameters in driving task performance. While a theoretical framework has been established for exploring the role of response thresholds in task allocation and division of labour (Loftus et al., 2021; Weidenmüller et al., 2019), there is an urgent need to incorporate different response parameters into this framework to obtain a comprehensive understanding of how social insect groups function.

## Data Accessibility

The NETLOGO code to run these simulations, the datasets obtained, and the Python and R codes used for the analysis are available from Zenodo via the DOI: 10.5281/zenodo.5217630.

## Author Contributions

Conceptualization: S.R., A.B., E.A.G.; Methodology: S.R., E.A.G.; Software: S.R., E.A.G.; Formal analysis: E.A.G.; Investigation: S.R., E.A.G.; Writing – original draft: S.R., A.B., E.A.G.; Writing – review & editing: S.R., A.B., E.A.G.; Supervision: A.B., E.A.G.; Funding acquisition: A.B.

