## Supplementary Information for "Environment dependent benefits of inter-individual variation in honey bee recruitment"

### Supplementary Tables

**Table S1**

| **Variable** | **Value** |
|---|---|
| Number of scouts | 30 |
| Number of recruits | 270 |
| Mean ± SD individual probability modulator of bees* | 0.255 ± 0.1 |
| Mean ± SD individual dance intensity modulator of bees* | 0.255 ± 0.1 |
| Time steps per day | 8640 |
| Probability that a scout will leave the nest | 0.815 |
| Probability that an idle recruit will scout | 0.00009 |
| Mean ± SD handling time on patch | 180±60 |
| Nectar handling time in nest | 6 |
| Distance of feeders from nest$^{+}$ | 80 patches |
| Nectar quality at feeders$^{+}$ | 0.8 M |
| Agent Speed | 0.7 patch width / time step |
| Levi flight parameter | 2.4 |
| Number of recruits following the dance of a bee | 1 |
| Probability of dance success | 0.25 |

The state variables used in the agent-based model. Variables marked with a (*) are individual specific variables unique to our model. Variables marked with a (+) have been modified from the original model for experiment 1. The other parameters in our model are similar to the model in Schürch and Grüter, 2014.

**Table S2**

| Contrast | Marginal Means [Confidence Interval] | Difference Estimate | z-ratio | p value |
|---|---|---|---|---|
| Model1 - Model2 | Model 1: 0.0099 [0.0081 - 0.0116]<br>Model 2: 0.1125 [0.1058 - 0.1193] | -0.103 | -29.233 | < .001 |
| Model1 - Model3 | Model 1: 0.0099 [0.0081 - 0.0116]<br>Model 3: 0.6365 [0.6261 - 0.6469] | -0.627 | -116.583 | < .001 |
| Model1 - Model4 | Model 1: 0.0099 [0.0081 - 0.0116]<br>Model 4: 0.4978 [0.4870 - 0.5086] | -0.488 | -87.472 | < .001 |
| Model2 - Model4 | Model 2: 0.1125 [0.1058 - 0.1193]<br>Model 4: 0.4978 [0.4870 - 0.5086] | -0.524 | -82.911 | < .001 |
| Model3 - Model2 | Model 3: 0.6365 [0.6261 - 0.6469]<br>Model 2: 0.1125 [0.1058 - 0.1193] | -0.385 | -59.334 | < .001 |
| Model3 - Model4 | Model 3: 0.6365 [0.6261 - 0.6469]<br>Model 4: 0.4978 [0.4870 - 0.5086] | 0.139 | 18.152 | < .001 |

Difference estimates for pairwise comparisons of the marginal means of repeatability estimates for all four models obtained from the beta regression analysis, along with the z-ratio and the p value (corrected for multiple comparisons using Tukey’s HSD) associated with the comparisons.

**Table S3**

| Parameter | Food Density | Contrast | Difference Estimate | t-ratio | p value |
|---|---|---|---|---|---|
| Mean Dance Time | Low | **Model1 - Model2** | 1.16 | 3.87 | < .001 |
| | | **Model1 - Model3** | 1.43 | 4.81 | < .001 |
| | | **Model1 - Model4** | 3 | 10.08 | < .001 |
| | | Model2 - Model3 | 0.28 | 0.93 | 0.788 |
| | | **Model2 - Model4** | 1.85 | 6.2 | < .001 |
| | | **Model3 - Model4** | 1.57 | 5.27 | < .001 |
| | Medium | Model1 - Model2 | 0.62 | 2.07 | 0.165 |
| | | Model1 - Model3 | -0.42 | -1.4 | 0.501 |
| | | **Model1 - Model4** | 2.32 | 7.77 | < .001 |
| | | **Model2 - Model3** | -1.03 | -3.46 | 0.003 |
| | | **Model2 - Model4** | 1.7 | 5.71 | < .001 |
| | | **Model3 - Model4** | 2.73 | 9.17 | < .001 |
| | High | **Model1 - Model2** | -0.92 | -3.08 | 0.011 |
| | | **Model1 - Model3** | -0.97 | -3.25 | 0.006 |
| | | **Model1 - Model4** | 2.27 | 7.63 | < .001 |
| | | Model2 - Model3 | -0.05 | -0.18 | 0.998 |
| | | **Model2 - Model4** | 3.19 | 10.71 | < .001 |
| | | **Model3 - Model4** | 3.24 | 10.88 | < .001 |
| Mean Yield | Low | Model1 - Model2 | -0.88 | -1.02 | 0.739 |
| | | Model1 - Model3 | 1.12 | 1.28 | 0.574 |
| | | Model1 - Model4 | 0.64 | 0.74 | 0.881 |
| | | Model2 - Model3 | 2 | 2.3 | 0.098 |
| | | Model2 - Model4 | 1.53 | 1.76 | 0.295 |
| | | Model3 - Model4 | -0.47 | -0.54 | 0.948 |
| | Medium | Model1 - Model2 | 0.24 | 0.28 | 0.992 |
| | | **Model1 - Model3** | -3.66 | -4.2 | < .001 |
| | | **Model1 - Model4** | -3.08 | -3.55 | 0.002 |
| | | **Model2 - Model3** | -3.9 | -4.48 | < .001 |
| | | **Model2 - Model4** | -3.33 | -3.83 | < .001 |
| | | Model3 - Model4 | 0.57 | 0.66 | 0.913 |
| | High | **Model1 - Model2** | -5.57 | -6.4 | < .001 |
| | | **Model1 - Model3** | -4.32 | -4.97 | < .001 |
| | | **Model1 - Model4** | -3.44 | -3.95 | < .001 |
| | | Model2 - Model3 | 1.25 | 1.43 | 0.478 |
| | | Model2 - Model4 | 2.13 | 2.45 | 0.068 |
| | | Model3 - Model4 | 0.89 | 1.02 | 0.738 |
| Median Foraging Distance | Low | **Model1 - Model2** | -0.05 | -3.54 | 0.002 |
| | | **Model1 - Model3** | -0.08 | -5.58 | < .001 |
| | | Model1 - Model4 | 0.001 | 0.08 | > .999 |
| | | Model2 - Model3 | -0.03 | -2.03 | 0.175 |
| | | **Model2 - Model4** | 0.05 | 3.62 | 0.002 |

| | | | | | |
|---|---|---|---|---|---|
| | | **Model3 - Model4** | 0.08 | 5.66 | < .001 |
| | Medium | Model1 - Model2 | 0.03 | 1.8 | 0.273 |
| | | Model1 - Model3 | -0.0008 | -0.06 | > .999 |
| | | Model1 - Model4 | -0.003 | -0.2 | 0.997 |
| | | Model2 - Model3 | -0.03 | -1.86 | 0.246 |
| | | Model2 - Model4 | -0.03 | -2 | 0.188 |
| | | Model3 - Model4 | -0.002 | -0.14 | > .999 |
| | High | Model1 - Model2 | 0.02 | 1.73 | 0.308 |
| | | Model1 - Model3 | 0.02 | 1.25 | 0.598 |
| | | Model1 - Model4 | -0.009 | -0.66 | 0.912 |
| | | Model2 - Model3 | -0.006 | -0.48 | 0.963 |
| | | Model2 - Model4 | -0.03 | -2.39 | 0.079 |
| | | Model3 - Model4 | -0.03 | -1.9 | 0.226 |
| Mean Search Time | Low | **Model1 - Model2** | -8.62 | -3.58 | 0.002 |
| | | **Model1 - Model3** | -12.35 | -5.12 | < .001 |
| | | Model1 - Model4 | -0.54 | -0.22 | 0.996 |
| | | Model2 - Model3 | -3.73 | -1.55 | 0.409 |
| | | **Model2 - Model4** | 8.09 | 3.35 | 0.004 |
| | | **Model3 - Model4** | 11.82 | 4.9 | < .001 |
| | Medium | Model1 - Model2 | 6.15 | 2.55 | 0.053 |
| | | Model1 - Model3 | 1.34 | 0.56 | 0.945 |
| | | Model1 - Model4 | -0.001 | -0.0005 | > .999 |
| | | Model2 - Model3 | -4.81 | -1.99 | 0.191 |
| | | Model2 - Model4 | -6.15 | -2.55 | 0.053 |
| | | Model3 - Model4 | -1.34 | -0.56 | 0.945 |
| | High | **Model1 - Model2** | 6.97 | 2.89 | 0.02 |
| | | Model1 - Model3 | 6.1 | 2.53 | 0.055 |
| | | Model1 - Model4 | 1.16 | 0.48 | 0.964 |
| | | Model2 - Model3 | -0.87 | -0.36 | 0.984 |
| | | Model2 - Model4 | -5.82 | -2.41 | 0.075 |
| | | Model3 - Model4 | -4.95 | -2.05 | 0.169 |

Difference estimates for pairwise comparisons of the marginal means of different parameters for all four models in each of the food density conditions obtained from generalized linear mixed effects model analysis, along with the t-ratio and the p value (corrected for multiple comparisons using Tukey's HSD) associated with the comparisons. Comparisons significantly different at the $p < 0.05$ level are highlighted in bold

### Supplementary Figures

**Figure S1**

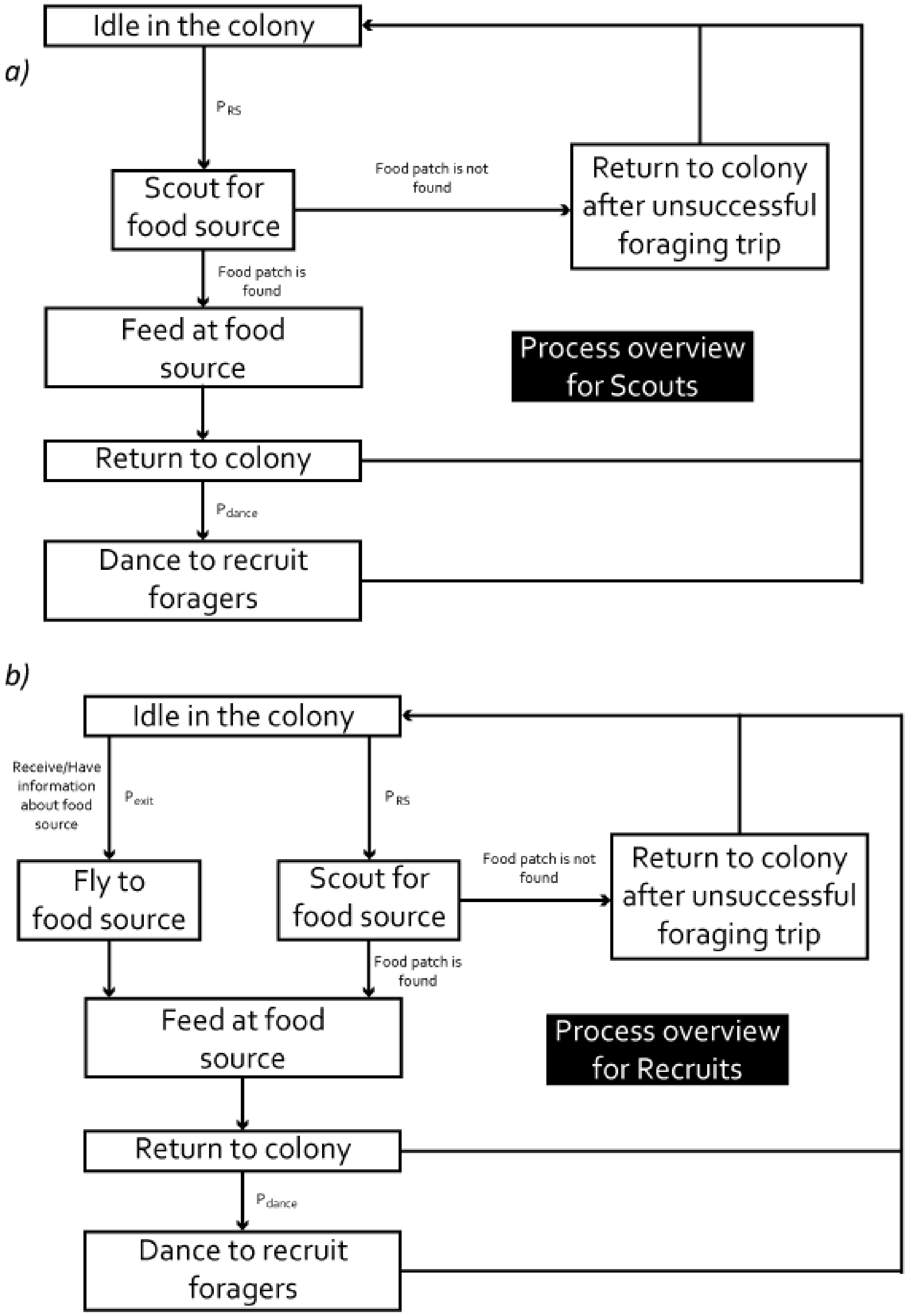


The process overview of (a) scouts and (b) recruits with the states that they can occupy at one time point in the model.

**Figure S2**

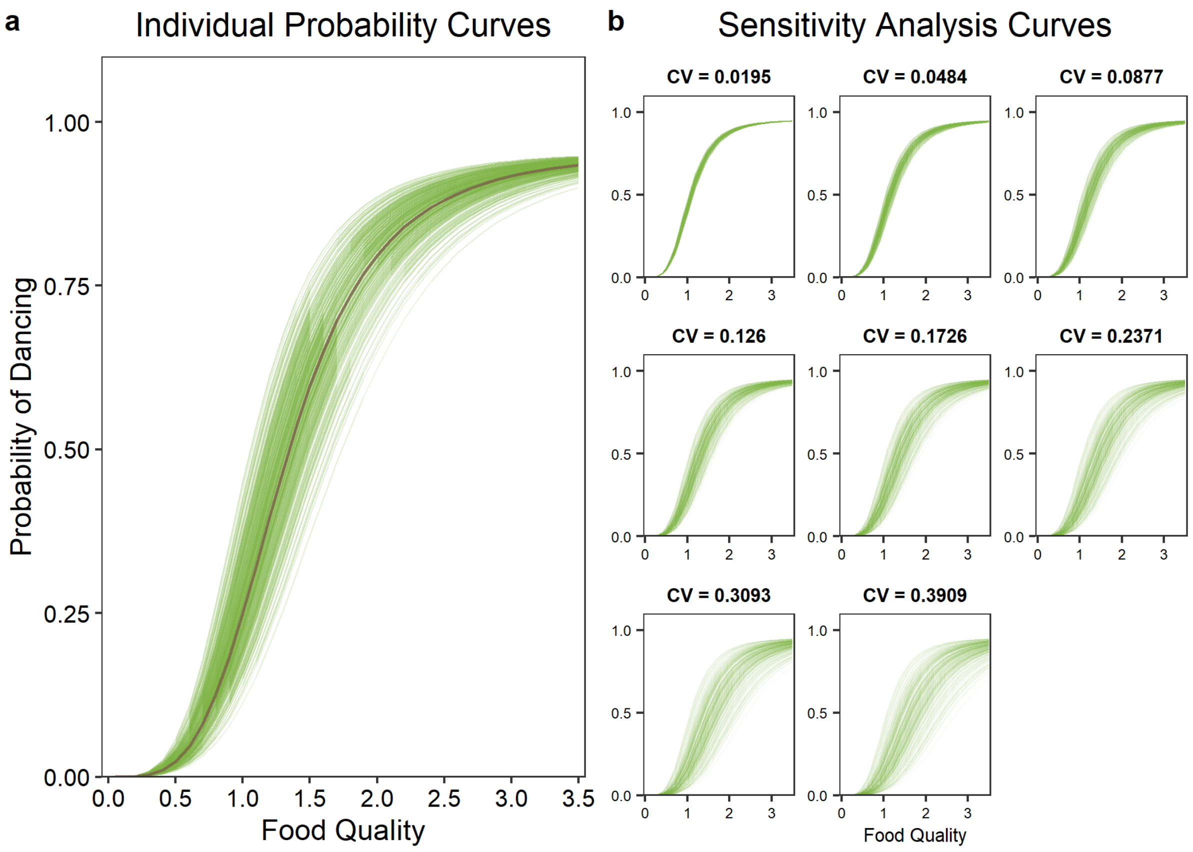


(a) The individual variation in the dance probability curves for 300 agents in one of the runs of Model 2 and 4. Each green curve corresponds to the relationship between probability of dancing and food quality for one of the agents, obtained from its probability modulator. The brown line corresponds to the relationship between probability of dancing and food quality for all agents in Model 1. (b) Individual variation in dance probability used in the sensitivity analysis. Each subplot corresponds to one particular coefficient of variation (CV) in the probability modulator. Green lines within each subplot represents the relationship between the probability of dancing and food quality for all 300 agents at this coefficient of variation.

**Figure S3**

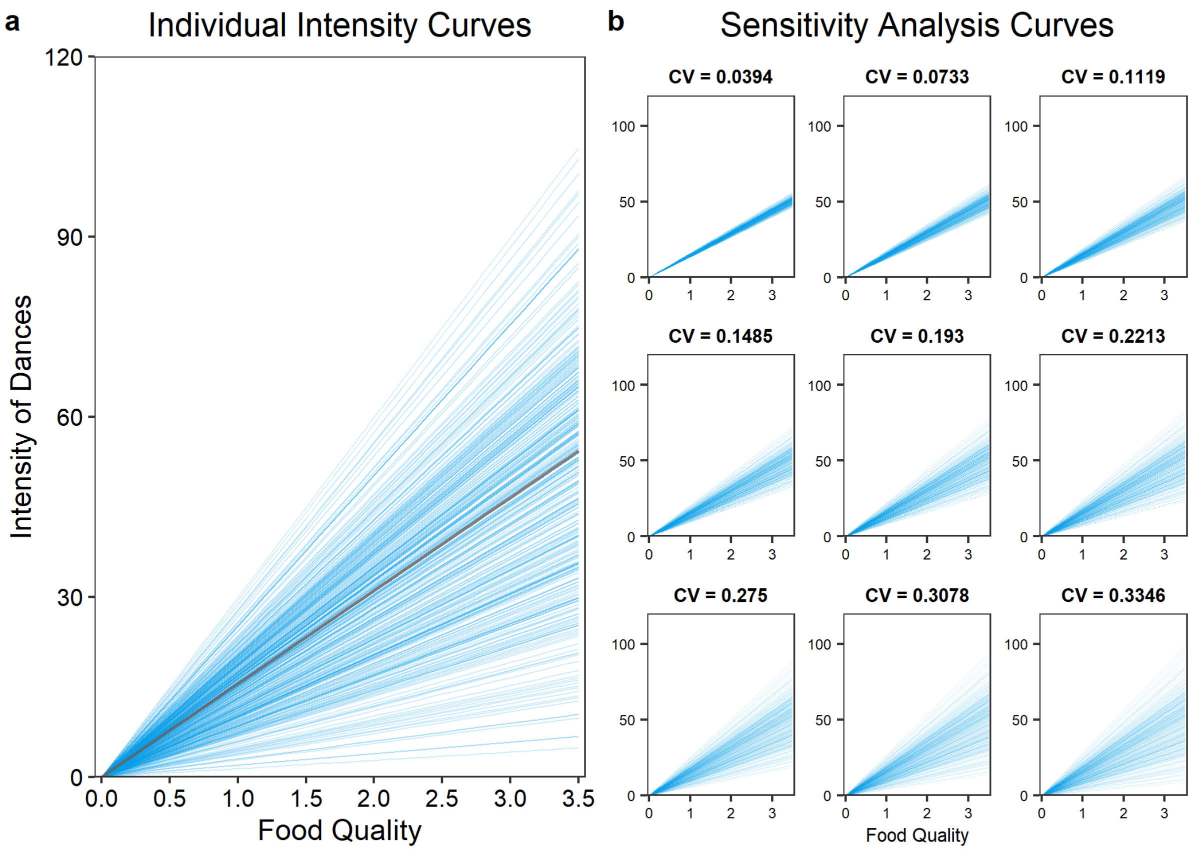


(a) The individual variation in the dance intensity curves for 300 agents in one of the runs of Model 3 and 4. Each blue curve corresponds to the relationship between intensity of dances and food quality for one of the agents, obtained from its intensity modulator. The brown line in the center corresponds to the relationship between intensity of dances and food quality for all agents in Model 1. (b) Individual variation in dance intensity used in the sensitivity analysis. Each subplot corresponds to one particular coefficient of variation (CV) in the intensity modulator. Blue lines within each subplot represents the relationship between the intensity of dances and food quality for all 300 agents at this coefficient of variation. The curves in (a) represent the variation in the intensity of dances at a coefficient of variation of 0.3705 of the intensity modulator.

**Figure S4**

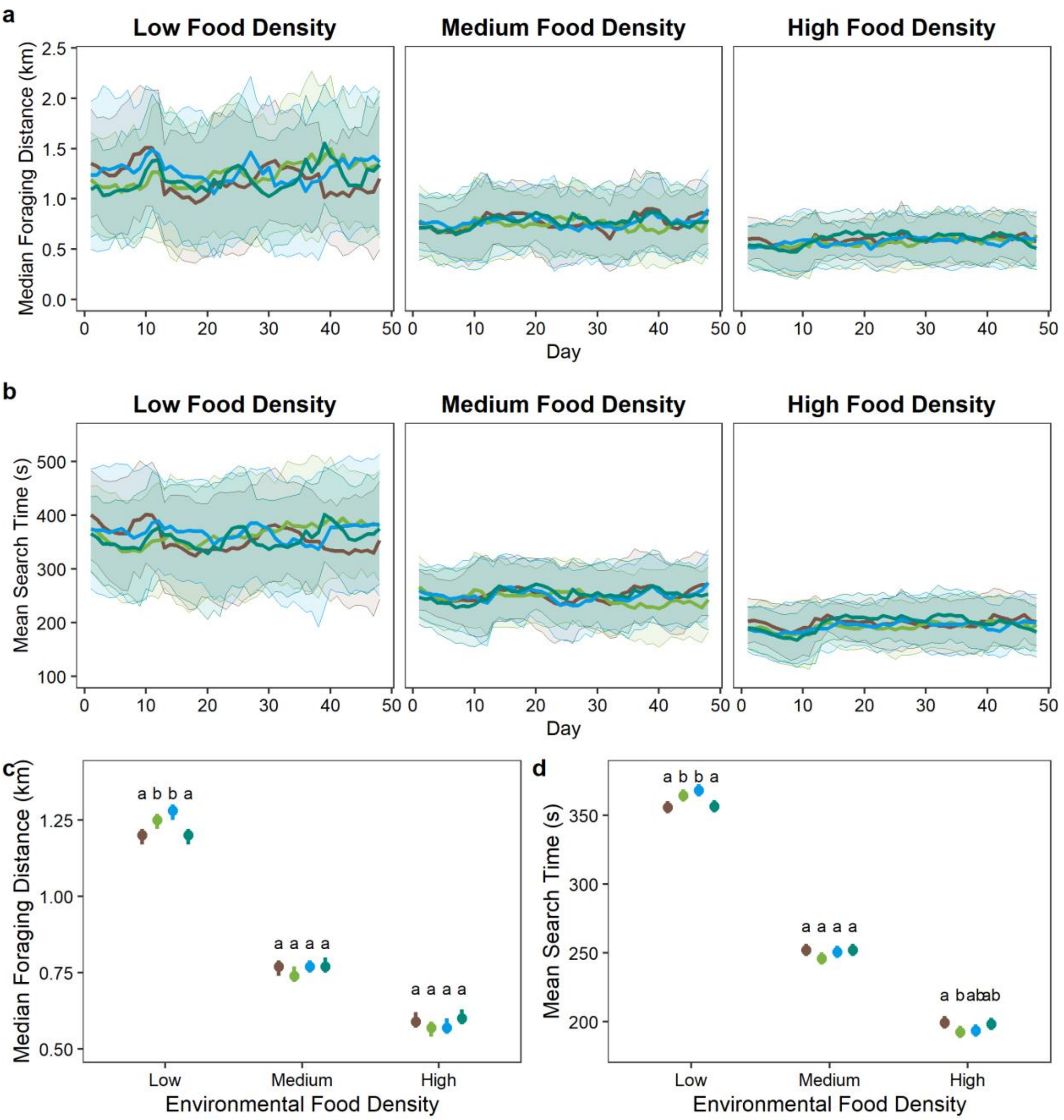


Effect of individual variation in recruitment behaviour on the colony's foraging activity under different environmental conditions. (a and b) Line plots showing the change in (a) median foraging distance in kilometres and (b) mean search time per forager in seconds throughout the simulations. The central line represents the mean value of all agents on each day from 40 simulations while the shaded region represents the standard deviation around this mean. (c and d) Comparison of the (c) median foraging distance and (d) mean search time, averaged over the whole simulation duration for all 40 simulations, across models and food density conditions. Circles represent the mean and error bars represent the standard deviation. The lines and shaded region in (a) and (b) and the circles and error bars in (c) and (d) are coloured according to the model: model 1 – brown, model 2 – green, model 3 – blue, model 4 – teal.
